# Rule-based habitat suitability modelling for the reintroduction of the grey wolf (*Canis lupus*) in Scotland

**DOI:** 10.1101/2022.03.01.482472

**Authors:** Vashti Gwynn, Elias Symeonakis

## Abstract

Though native to Scotland, the grey wolf (*Canis lupus)* was extirpated c.250 years ago as part of a global eradication drive. The global population has recently expanded, now occupying 67% of its former range. Evidence is growing that apex predators provide a range of ecological benefits, most stemming from the reduction of overgrazing by deer – something from which Scotland suffers. In this study, we build a rule-based habitat suitability model for wolves on the Scottish mainland. From existing literature, we identify the most important variables as land cover, prey density, road density and human density, and establish thresholds of suitability for each. Fuzzy membership functions are used to assign suitability values to each variable, followed by fuzzy overlay to combine all four: a novel approach to habitat suitability modelling for terrestrial mammals. Model sensitivity is tested for land cover and prey density, as these variables constitute a knowledge gap and an incomplete dataset, respectively. The Highlands and Grampian mountains emerge strongly and consistently as the most suitable areas, largely due to high negative covariance between prey density and road/human density. Sensitivity testing reveals the models are fairly robust to changes in prey density, but less robust to changes in the scoring of land cover, with the latter altering the distribution of land mainly through the 70 – 100% suitability range. However, in statistical significance tests, only the least and most generous versions of the model emerge as giving significantly different results. Depending on the version of the model, a contiguous area of between 10,139km^2^ and 18,857km^2^ is shown to be 80 to 100% suitable. This could be sufficient to support between 50 and 94 packs of four wolves, if the average pack range size is taken to be 200km^2^. We conclude that in terms of habitat availability, reintroduction should be feasible.

## Introduction

The grey wolf (*Canis lupus*) is native to Scotland, but was extirpated by humans c.250 years ago (1,2). This persecution was part of a global eradication effort that brought wolf numbers to their lowest point between the 1930s and 1960s (3). However, due to subsequent legal protection and conservation, the wolf population has expanded once again, and now occupies 67% of its former global range, including substantial expansion in mainland Europe (4). It is, therefore, unnecessary to re-establish wolf populations in the UK in order to conserve the species, but there is growing evidence that the presence of native apex predators brings with it a range of ecological benefits (2,4,5). Ripple et al. (4) showed that large carnivores are necessary to maintain biodiversity and ecosystem function, and that their roles cannot be fully reproduced by humans, and Atkins et al. (5) state that the elimination of large carnivores can suppress plant regeneration, due to population expansion and behaviour changes in herbivores. Moreover, the grey wolf can cause mesopredator cascades (affecting both mesopredators and their prey), and tri-trophic cascades (affecting every level of the food-web down to plants) (4). Such benefits are needed in Scotland, where deer densities are beyond ecological sustainability, and where red deer can reach a density of 150/km^2^ in some areas in winter (2,6). This has a serious impact on the structure, composition and function of Scottish ecosystems, especially on tree regeneration, through overgrazing and over-browsing (2,6). The 1995 reintroduction of grey wolves into Yellowstone National Park is considered instructive as to what may happen should wolves be reintroduced to the Scottish Highlands, as they share almost identical key species (grey wolves, elk/red deer, aspen) (2). In Yellowstone, just a few wolves have had profound effects, including tri-trophic cascades that ultimately improved river hydrology, and increased abundance and diversity in many species (2).

Nilsen et al. (1) predict that if wolves were present in Scotland for 60 years, deer densities would decline to 7/km^2^, with >50% reduction in some places. This is in line with the Deer Commission for Scotland’s target of 6/km^2^, and would greatly relieve the current financial burden of annual hind culling in pursuit of this target (1). Additionally, it is proposed that the re-establishment of the “Landscape of Fear” would produce behavioural changes in deer, and thus ecosystem benefits, beyond what reduction of numbers could achieve (2,7). Other benefits could include significant wolf-related tourism and carbon sequestration due to regenerating woodland (1,4).

While wolf reintroduction in Britain is not currently being considered, the government’s 25 Year Environment Plan sets out policy commitments to provide “opportunities for the reintroduction of native species” (8). Reintroductions and rewilding are currently popular, and the reintroduction of another keystone species – beavers – has received much attention and support. There is also growing emphasis on the ecological importance of intact ecosystems, e.g. see Plumptre et al. (9). In light of this, and the well-documented possible benefits of apex predators outlined above, the feasibility and desirability of wolf reintroduction in the UK needs to be assessed. Manning et al. (2) note the importance of a pre-existing body of research should reintroduction be considered in the future. This study is limited to Scotland because it is the area of the UK likely to be most suitable, due to its extensive deer-filled wild lands and low human density. Additionally, only the Scottish mainland was considered, as this is where any reintroduction programme would likely take place. Previous studies have explored some aspects of large predator reintroduction in Scotland, including modelling hypothetical impacts of wolves on the deer population (1,10), and mapping habitat and likely population expansion if lynx were reintroduced (11,12). Sandom et al. (10) take into account some habitat elements in their model of a hypothetical large fenced Highland reserve containing wolves. However – to our knowledge – no one has yet created a wolf habitat suitability model for all of mainland Scotland.

There are many existing predictive habitat suitability models for wolves in countries where they are already extant (notably in Italy, Switzerland, Poland, Germany and the northern USA) (13–22). Usually, the environmental characteristics of the areas in which wolves are already present are used to train the model (often a logistic regression model), which is then applied across the country or region to identify other areas that may be suitable. The situation in Scotland is fundamentally different, as wolves are not currently extant there, and neither Italy, Switzerland, Poland, Germany nor the northern USA can be considered sufficiently similar to Scotland, as regards land cover, climate and elevation, to be able to apply their habitat selection models directly. Fuzzy logic analysis is widely used in predictive modelling, including marine and aquatic habitat suitability modelling (23–26). It recognises marginal locations that sit on the boundaries of classes by assigning *likelihood* of class membership to each location (23,27)However, with the exception of the study by Zabihi Afratakhti et al. (27) fuzzy logic analysis is notably absent from the field of terrestrial habitat suitability modelling. Thus its use in this study represents a novel approach to mammalian habitat suitability modelling.

Here, we assess habitat suitability for wolves in mainland Scotland employing a rules-based approach, based on existing knowledge about wolf ecology (23), and fuzzy logic analysis, which allows for dataset inaccuracies and uncertainty in both the definition of attribute classes and in the measurement of the phenomenon. We carry out a sensitivity analysis of the model for the variable whose suitability is most uncertain (land cover), and for the variable for which we have the least data (prey density). Along with the use of fuzzy logic analysis, this allows for the incorporation of uncertainty in modelling and subsequent decision-making (28). This is particularly important in habitat suitability studies, as they contain numerous possible sources of error and/or uncertainty, e.g. spatial data inaccuracies, definition of rules based on other environments, etc (28,29).

## Methods

### Study Area

Scotland is a north-west European nation of 78,352km^2^, occupying the northern third of the island of Great Britain (Fig 1) (30,31). It has a temperate oceanic climate, that is wetter in the west with milder winters. Mean temperature in the coldest month is approximately 4°C, and in the warmest month, 14°C. Annual precipitation ranges from 635mm - >1000mm east to west, and significant snow falls on land above 460m in the winter (30). Glaciated in the Pleistocene, the Highlands in the north are mountainous and rugged, whereas the Central Lowlands are relatively flat, and the Southern Uplands are hilly (31). Almost all of Scotland’s primary forests have been cleared, and peatlands are widespread on the moors and hills, which are largely used for sheep farming and deer and grouse estates (30,31). Only 10% of the UK’s population live in Scotland, 75% of which dwell in the Central Lowlands, leaving rural areas sparsely populated (30).

**Fig 1:**
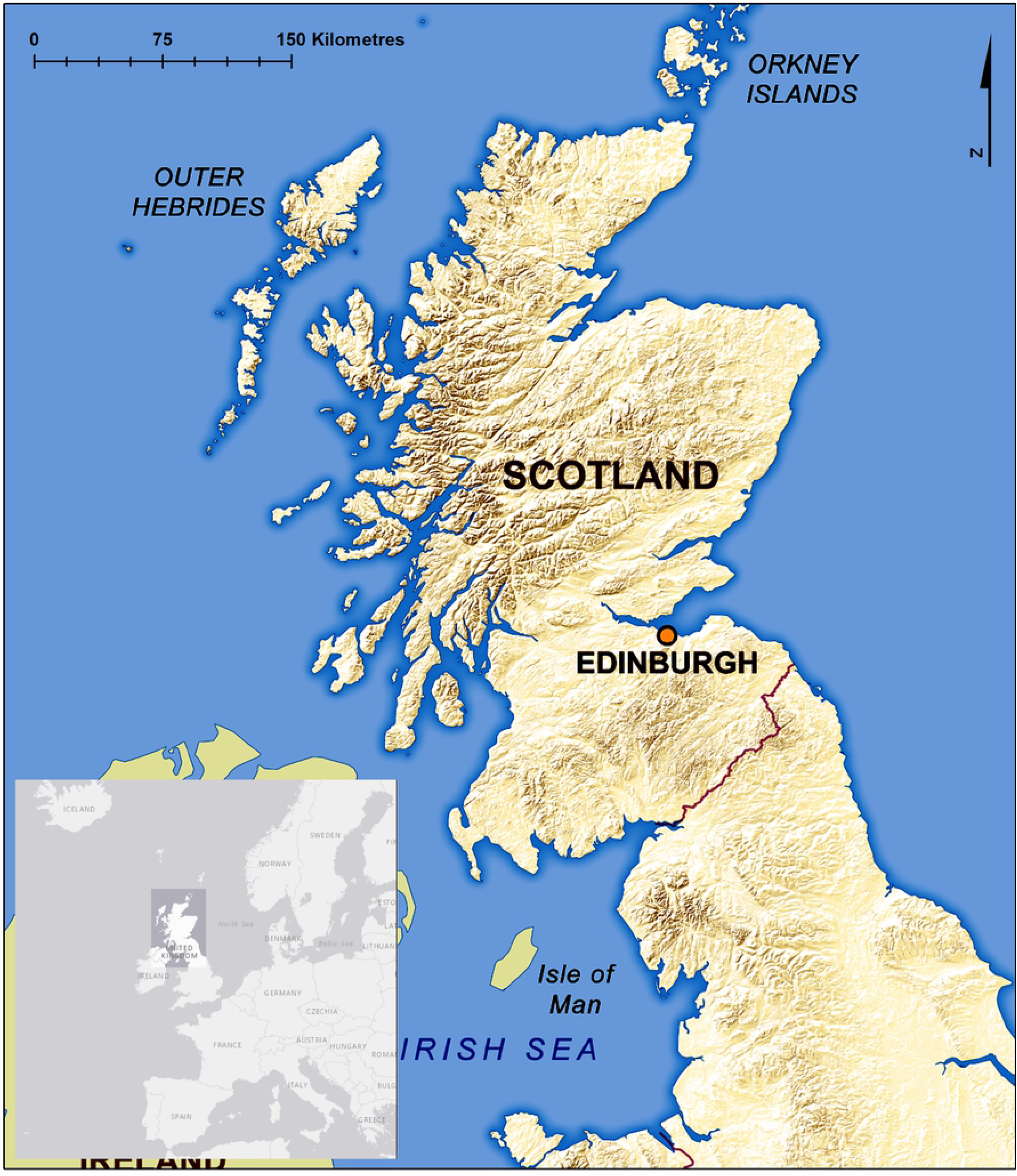
Physical map of Scotland and its location within north-west Europe inset. The mountains of the Highlands/Grampians in the North, the belt of the Central Lowlands, and the hills of the Southern Uplands can be clearly seen (Ordnance Survey, 2013; OpenStreetMap, n.d.).

### Rationale

Wolf habitat must be considered at a landscape scale, due to the size of the pack territories, which are typically 100 – 200km^2^, but vary greatly (13,17,32), and due to wolves’ long-distance dispersal, which can be hundreds of kilometres (32). Many European studies find there is a strong correlation between wolf presence and forest cover (14,15,19,20), but it must also be recognised that in these countries, areas with *low* human influence and *high* prey density tend to have correspondingly *high* forest cover. For instance, the Swiss Valais is 22% forest, and nearly all ungulates are restricted to forest habitats, especially in winter (18). Jędrzejewski et al. (19,20) also attribute Polish wolf pack preference for forest cover to avoidance of humans. This association between high forest cover and high prey density/low human presence is not the case in Scotland, which is only 18.5% forest (mostly conifer plantations) and where heathland and upland bog constitute the majority of its unpeopled and deer-stocked wild lands (33). Meanwhile, many American models find that wolf presence and abundance is more directly related to prey density or human-caused mortality risk than land cover (13,16,21). For instance, the wolf packs in the Canadian Arctic follow the caribou herds regardless of habitat (34). Similarly in north-eastern USA, prey availability and not habitat type explained 72% of spatial wolf population variation (16). Road density is also recognised as a crucial factor in habitat suitability in multiple studies (e.g. (35,36)). This difference between American and European studies suggests that either different limiting factors are at play, or that high covariance makes it hard to disentangle the importance of each variable.

Similarly, slightly different wolf predation behaviours emerge from different European studies, though wild ungulates always predominate (though see Ciucci et al. (14), for scavenging behaviour on garbage dumps) (32,37). Though roe and red deer form the majority of wolf diets in most European studies, some studies suggest red deer are preferred (though roe deer often still make up the majority of the diet due to higher availability) (19,38–42). Therefore, it is likely that in Scotland, both roe and red deer would be predated, though there may be a preference for red deer where they are available. Fallow and Sika deer are also found in Scotland, but little data exists for wolf predation on these species (43).

Despite these variations across studies, land cover, prey density, road density and human density emerge as the most important factors in wolf habitat suitability. As regards land cover, we noted which cover types are associated with wolf presence and absence, but as regards the other three variables – which are continuous rather than categorical variables – we needed to establish thresholds of suitability and unsuitability.

Prey densities that characterise areas of wolf presence (i.e. suitable habitat) vary across studies. Jędrzejewski et al. (20) noted a drop-off in wolf presence only when prey densities were as low as 0.6 deer/km^2^, but other studies find density requirements of at least 4/km^2^, with up to 13 elk per km^2^ recorded in the wolf ranges in Yellowstone National Park (10,21,22,37,44).

As regards roads, road density (km/km^2^) is the standard metric used in studies that assess wolf responses to roads (13,16,18–20,35,36). Recorded road densities in areas of wolf presence (i.e. suitable habitat) vary from 0.2km/km^2^ to around 0.4 or 0.5km/km^2^. Though there are areas with road densities of 0.7km/km^2^ being resettled by wolves, most studies thus far find such densities to be largely unsuitable (13,16,18,35,36,45).

Recorded human densities in areas of wolf presence (i.e. suitable habitat) begin at 0.43/km^2^ (21), but there is some variation in where the upper limit lies, with the same study finding an average of just 2.33 people/km^2^ in non-pack areas, whereas other studies record human densities all the way up to 36.7/km^2^, especially in Europe (13,18,22,45–47).

Given these varying preferences and behaviours in different regions (none of which are entirely comparable to Scotland), it may seem challenging to derive suitability rules that would apply to Scotland. However, the wolf’s generalist ecology helps to offset this. Wolves are not habitat specific, and nor are they necessarily wilderness species. They have colonised habitats throughout the northern hemisphere wherever they are protected from persecution, from 20° north up to the Arctic (3,13,48). Wolf core ranges have been found to include a wide range of habitats in addition to forests, including pasture, chaparral, eskers, heath tundra, and human garbage dumps (14,34,49). Recently wolves successfully recolonised a National Park in the Netherlands, an urbanised country with an average human population density of 512/km^2^ (against Scotland’s 70/km^2^) (50,51). Additionally, Scotland was until recently (ecologically speaking) part of the wolf’s range, and it was eradicated by persecution rather than by a lack of suitable habitat. Of course it may be fallacious to assume that because Scotland offered suitable wolf habitat in the past, it continues to do so now and in the future, but this only increases the need for rigorous study to test if that is so (29). Osborne and Seddon (29) recognise that it is essential to model extensively before reintroduction of any species, and that unsuitable habitat may be the main reason for reintroduction failures in the past.

Once the most important factors had been identified, and suitability thresholds established for each, fuzzy membership was applied to GIS datasets of the three continuous variables across the Scottish mainland, while land cover types were allocated scores and likewise mapped. The resulting output maps were combined using fuzzy overlay (Fig 2). This process was applied to six variations in input data, to explore uncertainty and to test sensitivity. The result is a set of six output maps, each containing all four variables, grading the Scottish mainland according to its suitability as wolf habitat.

**Fig 2:**
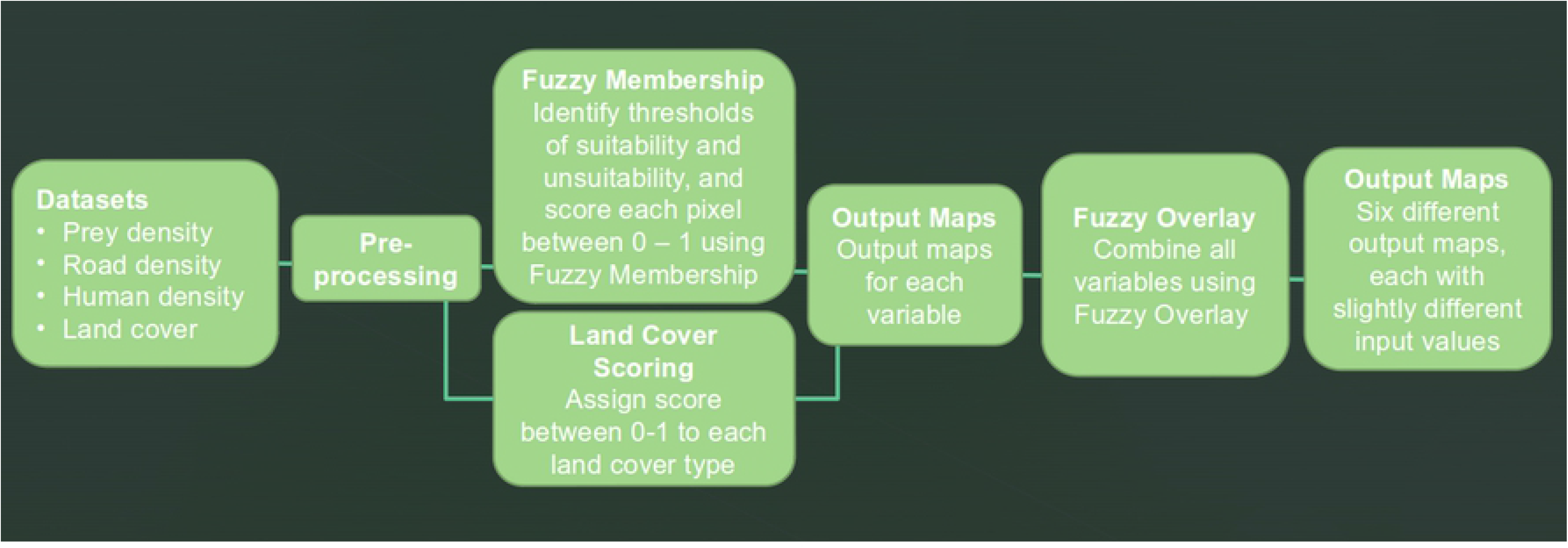
Flowchart of process. A summary of the analysis process, from input datasets to output maps. Note there are output maps for each variable individually, and then further output maps of all four variables combined using fuzzy overlay.

### Datasets

Spatial datasets for the four variables for Scotland were assembled in a GIS (ArcGIS Desktop 10.7.1 (52)), clipped to the Scottish mainland and converted to raster if originally in vector format (Table 1).

**Table 1.**
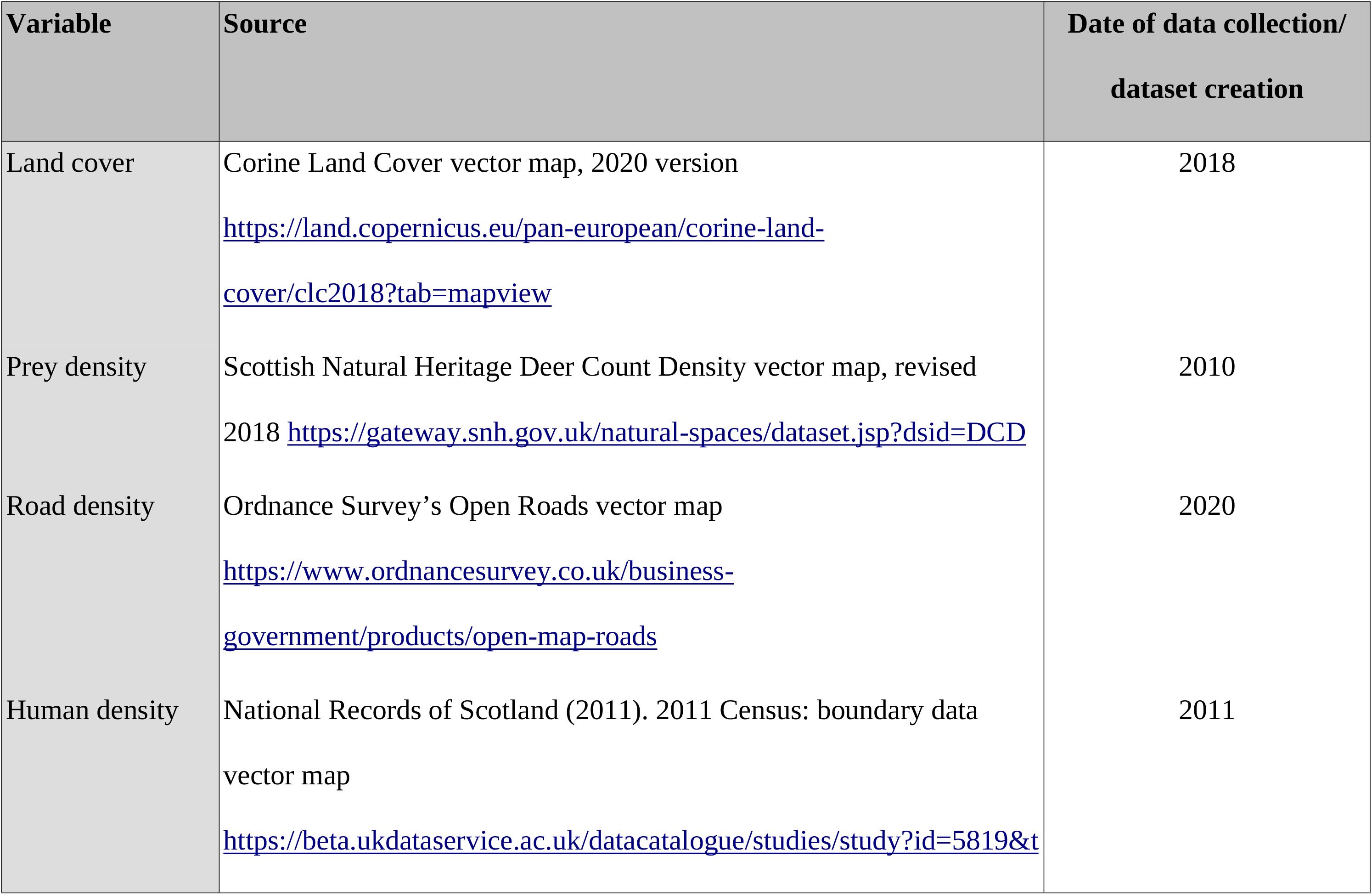
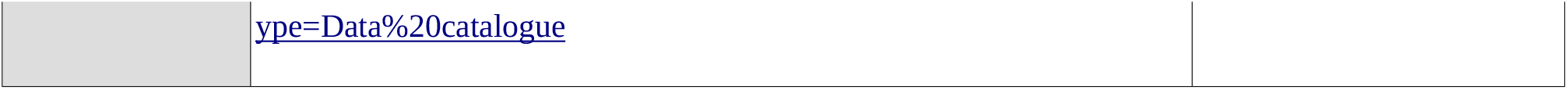
Datasets and their sources used in the analysis of habitat suitability.

| Variable | Source | Date of data collection/<br>dataset creation |
| --- | --- | --- |
| Land cover | Corine Land Cover vector map, 2020 version<br>https://land.copernicus.eu/pan-european/corine-land-cover/clc2018?tab=mapview | 2018 |
| Prey density | Scottish Natural Heritage Deer Count Density vector map, revised<br>2018 https://gateway.snh.gov.uk/natural-spaces/dataset.jsp?dsid=DCD | 2010 |
| Road density | Ordnance Survey's Open Roads vector map<br>https://www.ordnancesurvey.co.uk/business-government/products/open-map-roads | 2020 |
| Human density | National Records of Scotland (2011). 2011 Census: boundary data<br>vector map<br>https://beta.ukdataservice.ac.uk/datacatalogue/studies/study?id=5819&t | 2011 |

### Land Cover

The Corine Land Cover map is a European land cover inventory in 44 classes, based on Sentinel and Landsat imagery. It is classified at three levels of increasing thematic detail (e.g. the land cover type “Wetlands” is subclassified into “Coastal Wetlands” and “Inland Wetlands”, which are themselves subclassified into five further classes). The middle level of classification was used, as it had an appropriate level of thematic resolution. Each land cover type was scored for wolf suitability according to the available literature on wolf habitat preferences described in the Rationale (Table 2). A similar approach was used by Sandom et al. (10) in their modelling of a hypothetical fenced reserve in Scotland, but here an index between 0 and 1 was used, as that is comparable to the way fuzzy membership is allocated (where 0 indicates unsuitable habitat, and 1 suitable habitat).

**Table 2.** Land cover suitability scores.

| Land cover | Score |
| --- | --- |
| Arable land | 0 |
| Artificial, non-agricultural areas | 0 |
| Coastal wetlands | 0.4 |
| Forest | 1 |
| Heterogeneous agricultural areas | 0.2 |
| Industrial, commercial and transport units | 0 |
| Inland waters | 0 |
| Inland wetlands | 0.2, 0.4 & 0.6<sup>a</sup> |
| Marine waters | 0 |
| Mine, dump and construction sites | 0 |
| Open spaces with little or no vegetation | 0.2, 0.4 & 0.6<sup>a</sup> |
| Pastures | 0 |
| Permanent crops | 0 |
| Shrub and/or herbaceous vegetation associations | 0.2, 0.4 & 0.6<sup>a</sup> |
| Urban fabric | 0 |

*<sup>a</sup>These land cover types were scored three times due to uncertainty about their level of suitability.*

The suitability of the land cover types “Inland wetlands”, “Open spaces with little or no vegetation”, and “Shrub and/or herbaceous vegetation associations” was particularly hard to score, because these land cover types are uncommon in areas where other wolf habitat suitability studies have been performed, and thus their suitability is unclear. Their level of suitability is also crucial as they are dominant habitats in Scotland. Therefore, the model was run once with them scored at 0.2, once at 0.4, and once at 0.6, so that the model sensitivity to this particular variable could be explored. These scores can be considered to indicate substantially unsuitable, somewhat unsuitable and somewhat suitable habitat, respectively.

### Prey Density

The map of deer density had a resolution of 1km^2^, with count data attached to each 1km^2^ cell. Due to the herding behaviour of red deer in the Highlands, deer density was highly aggregated, i.e. one cell could contain dozens of deer while those around it contained none, reflecting where the herd happened to be on the day of the count. As this snapshot did not accurately reflect the realised spatial density of deer over time, kernel density estimation (KDE) was applied to each herd location, with an output cell size of 50m and a search radius of 1480m (representing the average home range of a red deer) (53,54). This “smooths” the density over a wider area, in recognition of the fact that the herd will move around its range, and so the entire range may be considered to offer prey (55).

Data from multiple studies suggest roe deer are an important component in wolf diets (see Rationale). Unfortunately, almost no roe deer are included in SNH’s Deer Count Density map, and no alternative roe deer density data was found. However, Campbell is cited as stating that roe deer density in the Highlands is 7.4/km^2^ and in the Southern Uplands 5.5/km^2^ (43). Therefore analysis was performed once with only the SNH dataset, and then a second time in which roe deer were additionally incorporated. They were incorporated at the densities mentioned above in every km^2^ cell for the Highlands and Southern Uplands (in the council areas of Highland, Dumfries and Galloway, and Scottish Borders, to be precise). Higher deer density suitability thresholds were used in the second version to account for the smaller body size of roe deer. The values used were, as with the red deer, guided by existing literature on roe deer densities in wolf territories (22). Due to the paucity of roe deer density data, this should be considered only as indicative of whether roe deer presence/absence might strongly change the outcome of the model, and more research is no doubt needed.

### Road Density

Road density (km/km^2^) was calculated from the Open Roads vector map at a resolution of 0.5km^2^. No weighting was applied to roads of different rank, as this does not seem to be common practice in wolf/road studies, but this could be worthy of further investigation.

### Human Density

Human density (people/km^2^) was calculated from the boundary census data vector map and converted to raster. With population data available only at census boundary scale, this is the dataset with the coarsest resolution. These boundaries are small in urban areas, where the population is high, but large in rural areas.

### Processing

Thresholds that represented completely suitable conditions (scored as 1) and completely unsuitable conditions (scored as 0) were established from the literature as regards deer, road and human densities (Table 3). Fuzzy membership with a linear membership type was then applied to each of these datasets accordingly, resulting in maps showing the suitability of that variable for wolves across mainland Scotland. This process was repeated for the deer density dataset with additional roe deer densities incorporated. The land cover dataset could not be processed using this method, because although the suitability scoring applied is numeric, it is categorical rather than continuous data, and therefore has no marginal cases (27).

**Table 3.** Thresholds of suitability.

| Variable | Suitable | Unsuitable |
| --- | --- | --- |
| Prey density without roe deer | $\geq 5.5/\text{km}^2$ | $\leq 1/\text{km}^2$ |
| Prey density with roe deer | $\geq 7/\text{km}^2$ | $\leq 3/\text{km}^2$ |
| Road density | $\leq 0.23\text{km}/\text{km}^2$ | $> 0.7\text{km}/\text{km}^2$ |
| Human density | $\leq 2/\text{km}^2$ | $\geq 37/\text{km}^2$ |

All four outputs were then assembled using fuzzy overlay with overlay type gamma of 0.9. Due to the paucity of fuzzy analysis in habitat suitability studies, there was no justification for using a different value, but this could be investigated further. Because of the two versions of deer density and three versions of land cover datasets in use, this resulted in six output maps. These maps each incorporate all four variables, but with some changes in input values in the deer density and land cover variables (Table 4).

**Table 4.** Versions of land cover dataset and prey density dataset used in each model.

|  | Inland wetlands, Open spaces with little or no vegetation, and Shrub and/or herbaceous vegetation |  |  |
| --- | --- | --- | --- |
|  | Suitability score = 0.2 | Suitability score = 0.4 | Suitability score = 0.6 |
| **Without roe deer** | Model 1<br>(Fig 3a) | Model 2 | Model 3<br>(Fig 3b) |
| **With roe deer** | Model 4<br>(Fig 3c) | Model 5 | Model 6<br>(Fig 3d) |

The fuzzy overlay maps were reclassified into 10 equal classes of suitability using ArcMap’s Reclassify tool. The distribution of pixels across the classes could then be used to calculate area and proportion of the Scottish mainland falling into each class, i.e. what proportion of land is 0-10% suitable, 10-20% suitable, and so on. Finally the Create Random Points tool and the Sample tool (resampling technique: nearest) were used to extract cell values from the same 500 randomly-generated points on each of the 6 fuzzy overlays, and a test for difference was performed in SPSS. As the data was strongly non-normal (Shapiro-Wilk: p=<0.001), Kruskal-Wallis with all pairwise comparisons was used.

## Results

Fig 3 shows the maps resulting from fuzzy membership analysis on land cover (Fig 3a, 3b, 3c), prey density (Fig 3d and 3e), road density (Fig 3f) and human density (Fig 3g). Land cover and prey density have three and two variations respectively, as per Table 4.

**Fig 3:**
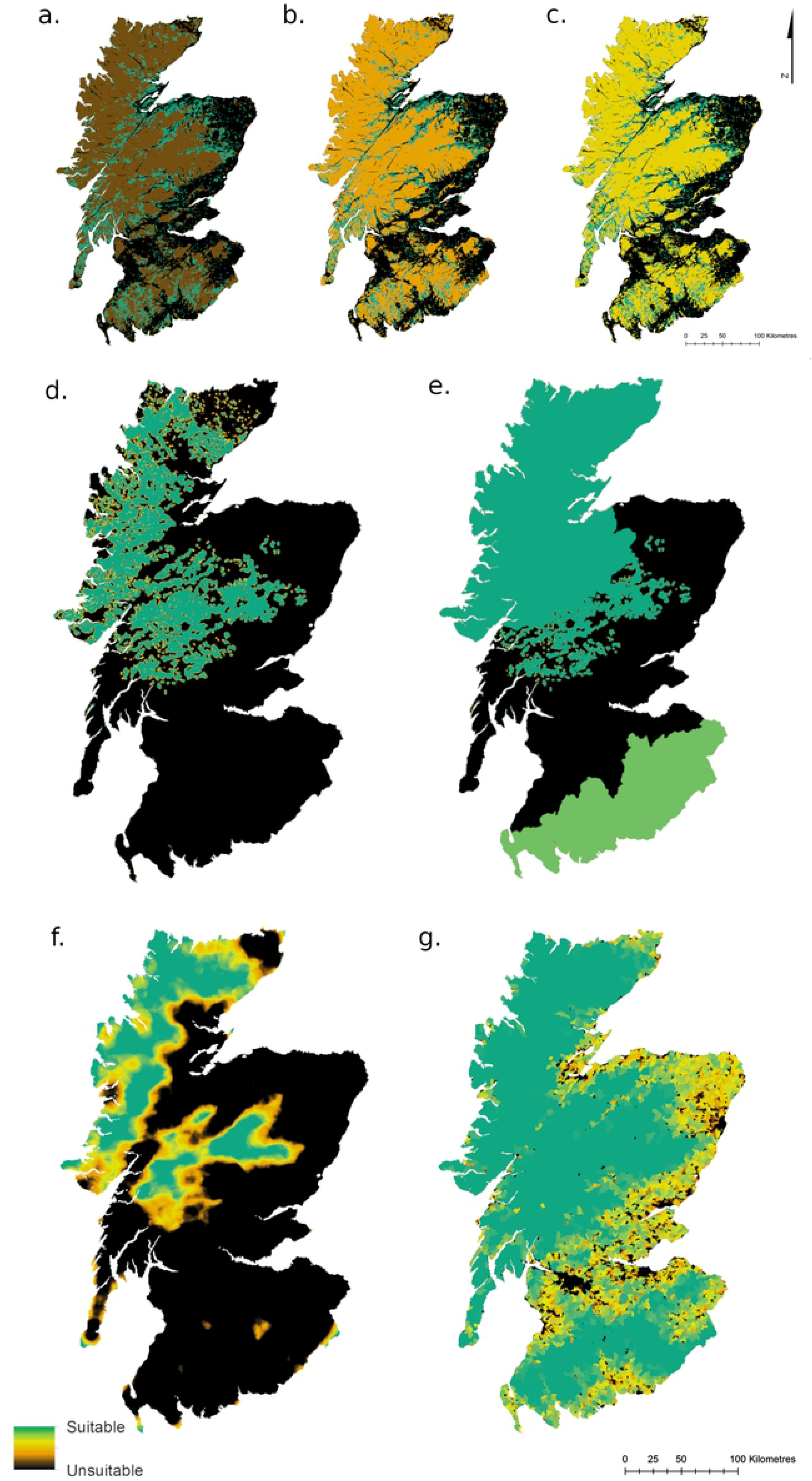
Fuzzy membership output maps. These show the suitability of the four variables: land cover (a,b,c) with three key types scored at 0.2, 0.4 and 0.6 respectively; prey density (d,e) without and with roe deer incorporated as a baseline density in two areas; road density (f) and human density (g). The Highlands and Grampians emerge consistently as the most suitable areas in all variables.

Fig 3a, 3b and 3c make explicit the large areas of Scotland covered by the three land cover types on which there is little suitability data: Inland wetlands, Open spaces with little or no vegetation, and Shrub and/or herbaceous vegetation associations. Meanwhile, it can be seen that red deer densities are largely suitable in the Highlands and Grampians (Fig 3d and 3e). The addition of a roe deer baseline density in three council areas increases the suitability of the Highlands and Southern Uplands. Human density (Fig 3g) is suitable for wolves across a large proportion of Scotland, but suitable road densities (Fig 3f) are far more limited, again to the Highlands and Grampians.

Fig 4 shows four of the six fuzzy overlay habitat suitability maps, each combining the four variables as per Table 4. Fig 4a and 4b do not include a roe deer baseline density along with red deer density, and Fig 4a and 4c score the three key land cover types at 0.2, whereas Fig 4b and 4d score them at 0.6. The two fuzzy overlay maps using the three land cover types scored at 0.4 can be seen in the Supporting Information (S1 Fig). The more suitable habitat is concentrated in the Highlands and Grampian Mountains in all model outputs.

**Fig 4:**
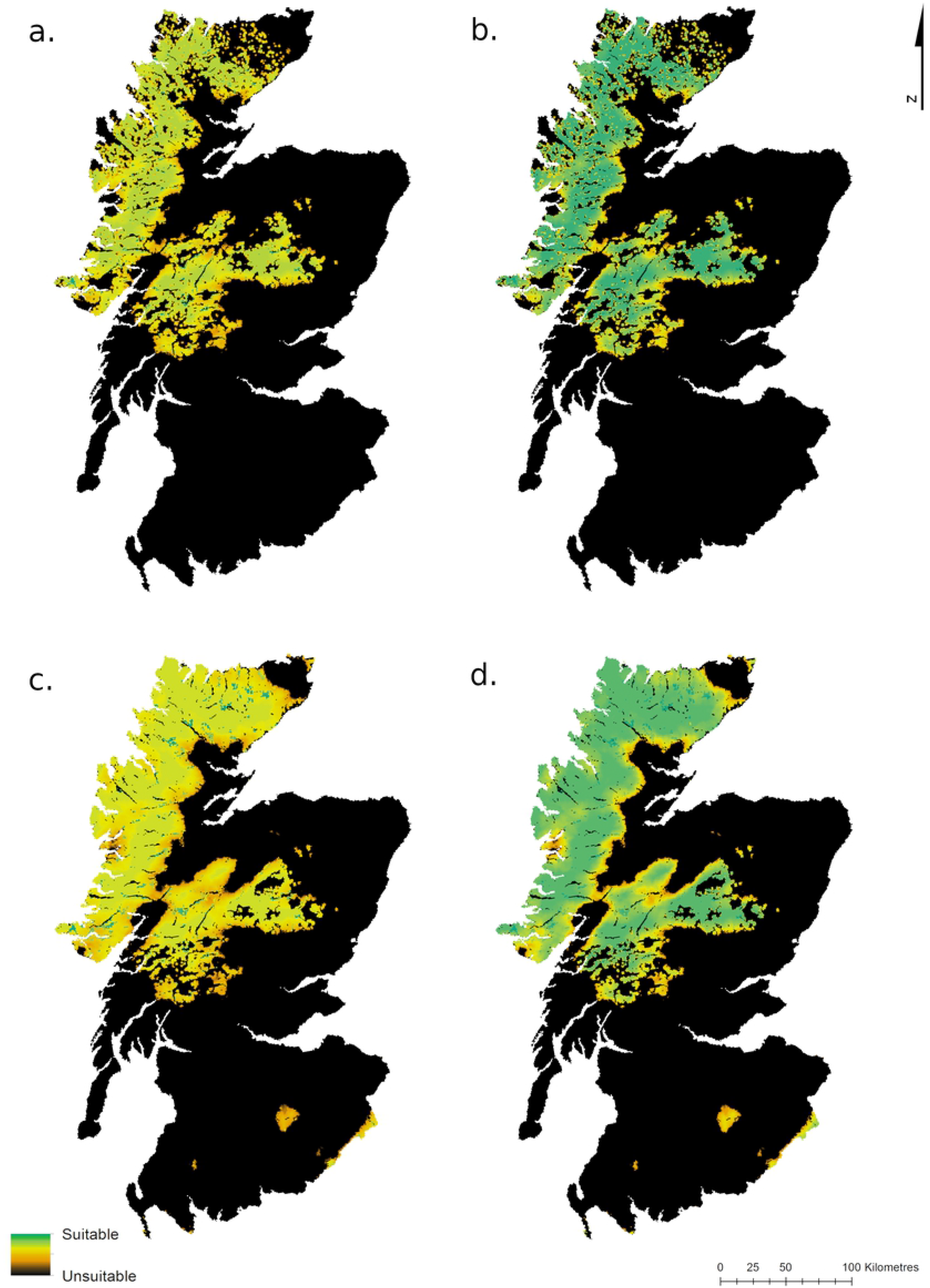
Overall wolf habitat suitability. Fig 4a corresponds to Model 1, Fig 4b to Model 3, Fig 4c to Model 4 and Fig 4d to Model 6 (Table 4).

The area and proportion of mainland Scotland falling into each of ten equal classes of suitability was calculated for each model (Table 5). This underlines the fact that the majority of Scotland is unsuitable according to these models, and that this does not vary much between models. There is more variation at the high suitability end of the scale, with between 0.6% and 21% of the area (or between 384km^2^ and 14259.5km^2^) rated most suitable (0.9 – 1.0), depending on the model used. Regardless of the model used, land is concentrated at either end of the scale of suitability, with very little semi-suitable habitat.

**Table 5.** Proportion of Scotland by suitability class.

| Suitability |  | Model 1 | Model 2 | Model 3 | Model 4 | Model 5 | Model 6 |
| --- | --- | --- | --- | --- | --- | --- | --- |
| **0.0 - 0.1** | Area (km<sup>2</sup>) | 51980.5 | 51981.5 | 51980.0 | 47305.5 | 47305.5 | 47306.0 |
|  | Proportion | 76.5 | 76.5 | 76.5 | 69.7 | 69.7 | 69.7 |
| **0.1 – 0.2** | Area (km<sup>2</sup>) | 0.0 | 0.0 | 0.0 | 0.0 | 0.0 | 0.0 |
|  | Proportion | 0.0 | 0.0 | 0.0 | 0.0 | 0.0 | 0.0 |
| **0.2 – 0.3** | Area (km<sup>2</sup>) | 1.0 | 0.5 | 0.5 | 0.0 | 0.0 | 0.0 |
|  | Proportion | 0.0 | 0.0 | 0.0 | 0.0 | 0.0 | 0.0 |
| **0.3 – 0.4** | Area (km<sup>2</sup>) | 15.5 | 12.0 | 6.5 | 9.5 | 5.5 | 4.0 |
|  | Proportion | 0.0 | 0.0 | 0.0 | 0.0 | 0.0 | 0.0 |
| **0.4 – 0.5** | Area (km<sup>2</sup>) | 86.5 | 46.0 | 37.0 | 58.0 | 30.0 | 20.0 |
|  | Proportion | 0.1 | 0.1 | 0.1 | 0.1 | 0.0 | 0.0 |
| **0.5 – 0.6** | Area (km<sup>2</sup>) | 346.0 | 200.0 | 146.0 | 225.0 | 139.5 | 101.0 |
|  | Proportion | 0.5 | 0.3 | 0.2 | 0.3 | 0.2 | 0.1 |
| **0.6 – 0.7** | Area (km<sup>2</sup>) | 1244.0 | 691.5 | 492.0 | 953.0 | 503.5 | 351.5 |
|  | Proportion | 1.8 | 1.0 | 0.7 | 1.4 | 0.7 | 0.5 |
| **0.7 – 0.8** | Area (km<sup>2</sup>) | 4097.5 | 2114.0 | 1492.0 | 4131.5 | 1917.5 | 1270.5 |
|  | Proportion | 6.0 | 3.1 | 2.2 | 6.1 | 2.8 | 1.9 |
| **0.8 – 0.9** | Area (km<sup>2</sup>) | 9755.5 | 7076.0 | 4237.0 | 14477.5 | 8820.5 | 4597.0 |
|  | Proportion | 14.4 | 10.4 | 6.2 | 21.3 | 13.0 | 6.8 |
| **0.9 – 1.0** | Area (km<sup>2</sup>) | 384.0 | 5791.5 | 9519.0 | 750.0 | 9190.0 | 14259.5 |
|  | Proportion | 0.6 | 8.5 | 14.0 | 1.1 | 13.5 | 21.0 |

Plotting this in graph form makes it plain that the different models have little impact on the distribution of land within the less suitable categories, but a larger impact on distribution in the more suitable categories (Fig 5 – note the logarithmic scale on the y-axis). It also becomes evident that the changes in land cover scoring have more of an impact on results than the addition of roe deer baseline densities.

**Fig 5:**
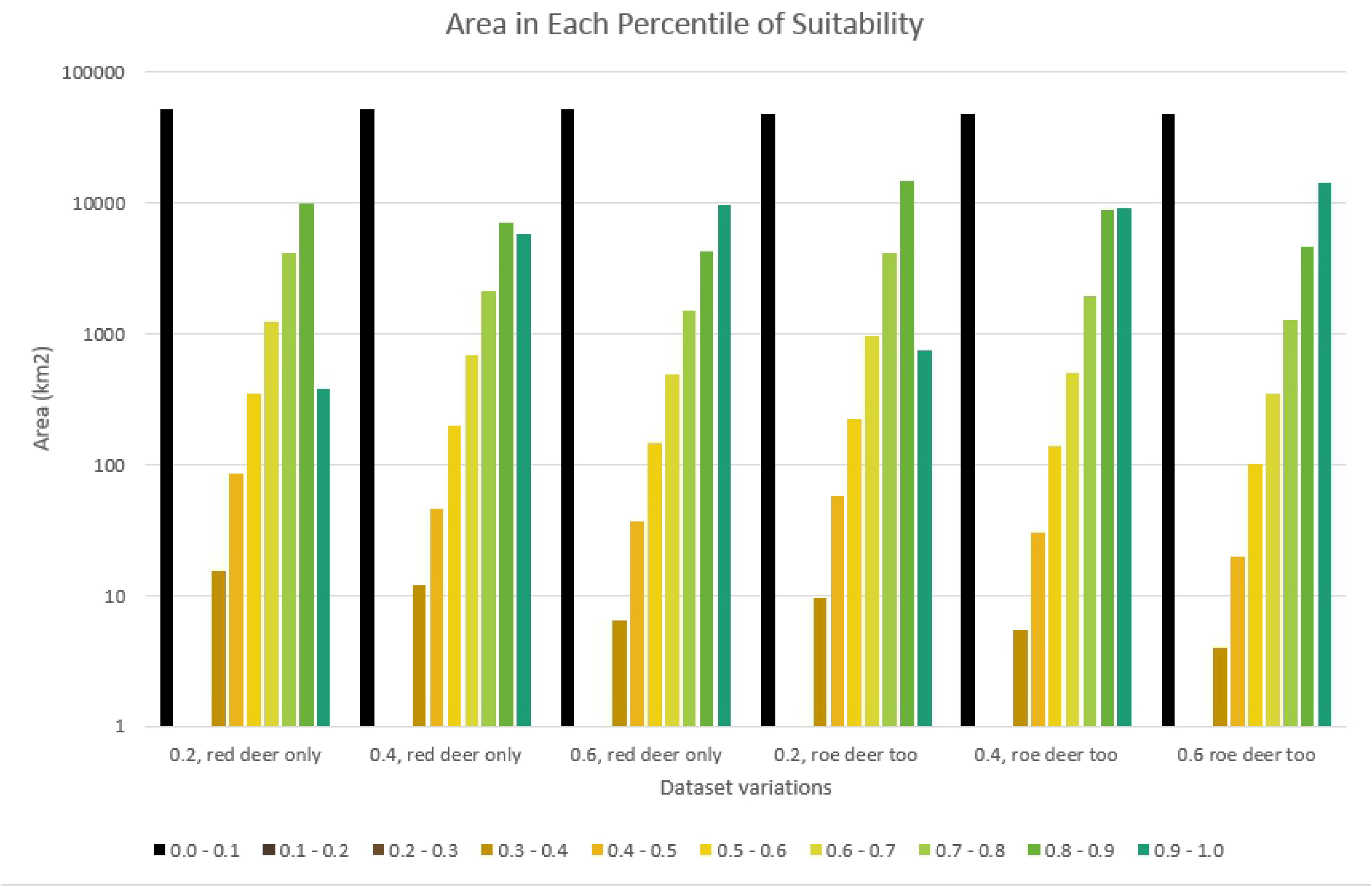
Area of Scotland by suitability class. Area of the Scottish mainland that falls into each of ten classes of suitability under different models (models 1 - 6 from left to right). Note that the Y-axis uses a logarithmic scale. All models show a similar pattern, with the distribution being highly uneven, and most areas being either completely unsuitable or substantially/completely suitable.

Testing for difference in a sample of 500 randomly-generated cell values taken at the same points for all six fuzzy overlay maps shows that there is only a significant difference in results between Model 1 and Model 6, i.e. the most and least “generous” models (H = −3.47, df = 5, p = 0.008, using the adjusted significance value).

## Discussion

Our results have shown that there is a high level of covariance between three of the variables, with the most suitable areas in terms of prey density, road density and human density all concentrated in the same regions (Fig 3). This results in the Highlands and Grampian mountains emerging strongly and consistently in Fig 4 as the areas most suitable for wolves in mainland Scotland. Though this area is contiguous, it is bisected by the A82 and the many lochs of the Great Glen, which could be barriers to wolf movement. Human density, prey density and road density are suitable throughout this region, and the addition of roe deer to the prey density map makes little difference. This is partly because in the Highlands the high densities of red deer already reach the suitability threshold, whereas in the Southern Uplands, the majority of the region is excluded anyway due to high road densities. It is also partly because the adjustment of the suitability thresholds upwards to account for the smaller body size of roe deer (Table 3) somewhat negates the gains of including them.

However, the suitability of the fourth variable, i.e., land cover, depends heavily on how suitable open heath and bog habitat is for wolves. It is the scoring of three land cover types in this variable that make the largest difference in the fuzzy overlay maps. Though in all cases the Highlands and Grampians still emerge as most suitable, differences in scoring mean they may be anywhere between somewhat suitable and completely suitable (between 0.7 and 1.0).

In terms of sensitivity, it can thus be concluded that the model is not particularly sensitive to the changes in prey density used here. However, it is somewhat sensitive to changes in land cover scoring: though the regions with highest suitability do not change, their level of suitability does.

As regards prey density, road density, and human density, the models could be considered relatively conservative. This is due to two reasons: the estimates of suitability and unsuitability adopted as thresholds were conservative; and prey density is likely to be higher and more widespread than the SNH’s deer counts suggest, as these counts mostly only include red deer spotted in open areas where and when a count is carried out. The British Deer Society’s distribution survey (56) finds that red deer are extant also in the Southern Uplands (which is not represented in the SNH counts), roe deer are common across Scotland, and fallow and sika deer are also found patchily in the Cairngorms, Highlands, west Scotland and the central Southern Uplands. However, it must be noted that regardless of prey density much of Scotland would remain unsuitable for wolves due to high road densities (Fig 3f).

Deer densities far exceed the threshold of suitability in much of the Highlands and Grampians with many areas holding >35/km^2^ according to SNH’s deer counts (after KDE processing). Hetherington and Gorman (43) quote an average density of 12.2 deer of all species/km^2^ in the Highlands, while 11–12/km^2^ is estimated by Sandom et al. (10). This is significant because wolf pack range size is largely determined by prey availability (16,32,37). Recorded range sizes vary from 33km^2^ to 6272km^2^ but average 100 – 200km^2^ (13,17,32,45). Fuller (37) found that at a density of 6.2 white-tailed deer per km^2^ (roughly half the Highland deer density) pack range size was only 116km^2^. Meanwhile, Sandom et al.’s (10) modelling suggested that a Highland fenced reserve of 600km^2^ would sustain 2 packs of 4 wolves for at least 100 years. At their most conservative, the models used in this study place an area of 10,139km^2^ between 80% and 100% suitable, and at their most generous, 18,857km^2^ is 80 – 100% suitable. This suggests that there may be sufficient wolf-suitable area to support between 50 and 94 packs of 4 wolves, if pack territory size is taken to be 200km^2^. However, it should be noted that the minimum size required for a single pack is unclear due to variations in range size recorded in the literature, and Sandom et al. (10) found that a fenced reserve of 200km^2^ was too small to support a Highland wolf pack for 100 years (though fencing brings with it implications that an unfenced population would not face). Additionally, a single – or even several – packs is not a self-sustaining population, as evidenced by the isolated wolf population on 544km^2^ Isle Royale in Lake Superior, whose numbers dwindled from 50 in 1980 to 2 in 2016 before reintroductions bolstered them (57). Indeed in 1992, the US Fish and Wildlife Service stated that an area of 25,000km^2^ was required for a self-sustaining wolf population (16).

However, the evidence base of what wolves require is still developing. Linnell et al. (48), Mech (3), and Mladenoff et al. (13) all note that with protection from persecution, wolves are recolonising areas previously thought unsuitable due to high human and road densities. Future observations of recolonising wolves in Europe are likely to be instructive as to what wolves prefer and tolerate. Clarification on wolf preferences as regards open upland habitats is particularly needed for assessment of Scottish habitat suitability, as studies currently conflict on how essential forests are for wolves (see Rationale). Though this is the largest data gap as regards modelling Scottish habitat suitability, other areas of future research could also include wolf response to roads. The standard in wolf habitat studies is employing a road density measure of km/km^2^. However, Jędrzejewski et al. (20) found that Polish wolves avoided a 250m wide belt along roads, so the use of buffer zones in models may be more beneficial. Additionally, there seems to be little research on the implications of roads of different class, and this could be explored further. Further research into Scottish habitat suitability would also benefit from more comprehensive deer density datasets, or else modelling of deer populations across Scotland that is more sophisticated than the KDE smoothing used here. Reinecke et al. (55) point out that one of the weaknesses of KDE is that it includes invalid areas (for instance a loch), and suggest minimum convex polygons or α-local convex hulls as alternative methods for modelling red deer. Dasymetric interpolation may also provide a more realistic model of deer densities (58). However, as our sensitivity testing indicated that changes in deer density did not particularly affect the model output, we did not refine our processing for this study.

There are many considerations regarding the return of wolves to Scotland that are beyond the scope of this study. These include the requirements of maintaining wolf genetic diversity and metapopulation, which would require either an area far bigger than that needed to support a few packs, or else regular introductions of additional animals. There are also implications arising from wolf dispersal (which can be many hundreds of kilometres) and social ecology (which is complex and could be negatively affected in a small, constrained population) (32). These implications are not explored here, but would be worth further study. While this study finds deer densities are easily sufficient to support wolves in the Highlands and Grampians, it does not model the long-term predator-prey relationship or the likely impact on deer population dynamics. Lastly, one of the most important considerations in any reintroduction project is public attitude and impacts on local communities. With apex predators there is a particular risk of human-wildlife conflict, especially due to livestock predation. Wolf reintroduction also has the potential to affect the recreational value of Scotland’s wild lands, and even its Protected Areas. See Sandom et al. (10) for modelling of predator-prey and population dynamics of wolves in the Highlands, and for discussion of the implications for recreation and Protected Areas. See Nilsen et al. (1) for modelling of predator-prey dynamics and ecological impacts should wolves be released in Scotland, and discussion of public attitudes and perceptions.

With the recolonisation of Europe by wolves, Britain increasingly becomes an outlier in its lack of apex predators. If the nation were not an island, wolves would likely soon cross our borders, if they had not already done so. As it is, short of escape from captivity, it is impossible for wolves to recolonise naturally, regardless of the suitability of our habitats or the desirability of their presence. Whether they return to Britain is a decision we must make actively, and in full consideration of the wolf’s requirements and impacts. Therefore conservationists need to anticipate evidence needs now (2). As well as the need to fill knowledge gaps (some of which are identified above) both Manning et al. (2) and Sandom et al. (10) advocate reintroduction experiments in large-scale Highland enclosures, which would allow us to discover the impacts of wolves on deer and ecosystems in general, in the context of the Highlands. This study supports previous conclusions that in terms of habitat suitability, the Highlands or Grampians would be the most appropriate places for wolf reintroduction. We recommend further research into the knowledge gaps outlined, and beyond that – should Scotland still appear suitable for wolves, as we find – consideration be made of the desirability of reintroduction.

## Conclusion

This study set out to identify the level of habitat suitability for wolves in the Scottish mainland. We have established thresholds of suitability and unsuitability from the literature as regards the four most important habitat variables: land cover, prey density, road density and human density. We mapped each variable according to its suitability across mainland Scotland using fuzzy membership analysis, and then combined all the variables into maps of overall suitability using fuzzy overlay. The Highlands and the Grampians emerged strongly as the most suitable areas, and sensitivity testing showed the model was fairly robust to changes in the prey density inputs, but less robust to changes in the land cover inputs, which we identify as an area in need of further research. Between 10,139km^2^ and 18,857km^2^ are found to be 80 – 100% suitable, depending on the model used, which may be sufficient to support between 50 and 94 packs of 4 wolves.

## 8. Supporting information

***S1 Fig: Habitat suitability maps produced using the three land cover types scored at 0.4.*** *These were produced by Model 2 (a) and Model 5 (b) in Table 4, respectively.*

